# *Shank3* mutation disrupts affective touch encoding in the dorsal medial prefrontal cortex of Beagle dogs

**DOI:** 10.64898/2026.09.01.748311

**Authors:** Yanhe Zhou, Yue He, Minna Dan, Lintao Jia, Kun Guo, Ying Fang, Li Hu, Xiang Yu, Jinfen Wang, Dajun Xing, Yong Q. Zhang

## Abstract

Individuals with autism spectrum disorder (ASD) often show aversion to affective touch (AT). However, the neural mechanism of this abnormality in cortices remains poorly understood probably due to the lack of effective animal models. Here, we used a canine model to address this issue by leveraging the intimate dog-human interactions. In a newly-designed heterospecific AT paradigm, we found that dogs carrying mutations in *Shank3*, a high-risk gene for ASD, avoided human AT. *In vivo* single-unit recording analysis showed that AT-evoked oscillations in the dorsal medial prefrontal cortex (dmPFC) were significantly altered in *Shank3* mutant dogs. *Shank3* mutation also reduced the number of neurons encoding AT in the dmPFC. Importantly, the aversion to AT and altered neural processing in *Shank3* mutants were largely rescued by a GABA_A_ receptor antagonist pentylenetetrazole. Together, these findings provide neural mechanisms for abnormal AT processing in ASD and suggest potential biomarkers for therapeutic strategies.

## Introduction

The sense of touch is critical to our interaction with the surrounding world and with each other (Moehring et al., 2018). Affective touch (AT), also known as social touch, conveys hedonic information that promotes affiliative behaviors and contributes to the well-being of social animals (McGlone et al., 2014; Morrison et al., 2010). AT, including behaviors such as cuddling, caressing, and hugging, is highly evolutionarily conserved across different species (Hertenstein et al., 2006). In nonhuman primates, rodents, and birds, allogrooming behavior is important for strengthening and maintaining social bonding, reciprocity, and hierarchy (de Waal and Suchak, 2010; Dunbar, 2010). AT also occurs in cross-species dyads such as humans and pets, especially domesticated cats and dogs.

A retrospective video analysis indicates that infants later diagnosed with autism spectrum disorder (ASD) often display social touch aversion as early as 9–12 months of age (Baranek, 1999). Subsequent studies confirmed that children’s early atypical sensory responses, such as the avoidance of touch, are predictive of later ASD (Mammen et al., 2015; Thye et al., 2018). Up to 90% of individuals with ASD have been reported to have alterations in sensory processing (Suarez, 2012). Of these, a subset display aversive behaviors including struggling, pulling, pushing, and gazing away to AT (Mammen et al., 2015; Tomchek and Dunn, 2007; Wiggins et al., 2009). In light of the growing body of clinical studies suggesting close association between sensory abnormalities and autistic behaviors, hyper- or hypo-sensory responses have been included in the diagnostic criteria for ASD. Imaging studies in ASD patients showed that the posterior insula, medial prefrontal cortex (mPFC), anterior cingulate cortex (ACC), and superior temporal sulcus (STS) are involved in the perception and processing of AT (Gordon et al., 2013; Voos et al., 2013). However, how AT is processed in the high-order cortical regions at neuronal level remains largely unknown.

Previous circuit studies on AT were mostly carried out in wild-type rodents focusing on subcortical nuclei (Hayashi et al., 2025; Yu et al., 2022). Yu et al. identified a dipeptidergic pathway from Tac1^+^ neurons in the lateral/ventral lateral periaqueductal gray (l/vlPAG) to the paraventricular nuclei (PVN) oxytocin neurons, through which pleasant touch experience promotes social interactions in mice (Yu et al., 2022). Similarly, an oxytocin pathway from the supraoptic nucleus (SON) to the ventromedial hypothalamus (VMH) facilitates human touch-induced play behaviors in rats (Hayashi et al., 2025). However, it is still unclear how high-order cortical regions process AT-associated tactile information.

Given that AT is evolutionarily conserved, and can occur between humans and pets, interaction between humans and dogs is an appealing model for investigating the cortical mechanisms underlying AT. Dogs have acquired advanced emotional and cognitive capacities during a long history of domestication (Range et al., 2009), show strong affiliation with humans (Hare and Tomasello, 2005), and have developed a rich repertoire of body language to express emotional states (Siniscalchi et al., 2018). These characteristics of domesticated dogs allow for quantitative behavioral measurements in controlled interaction settings with humans. Moreover, the ASD-related *Shank3* mutant Beagle dogs, displaying specific deficits in social interactions (Tian et al., 2023), provide an unprecedented and irreplaceable model for studying cortical processing of AT in a neuropathological condition.

Here, we take advantage of the dog models to investigate the neural mechanisms of AT using behavioral assays and electrophysiological analyses. In the human to dog AT paradigm, *Shank3* mutant dogs showed significantly elevated stress responses to gentle hand stroke, whereas wild-type (WT) dogs enjoyed it. *In vivo* single-unit recording analysis revealed that the dorsal medial prefrontal cortex (dmPFC) encodes both pleasant and unpleasant tactile information. We further observed that loss of *Shank3* disrupted AT encoding in individual neurons and led to altered oscillations in the dmPFC. Importantly, the GABA_A_ receptor antagonist pentylenetetrazole (PTZ) rescued the impaired neural encoding of and aversion to AT in *Shank3* mutants. Together, these results uncover the neural mechanism by which dmPFC processes AT with positive valence, and shed light on neural deficits that underlie AT aversion in ASD.

## Results

### Aversion to affective hand stroke in *Shank3* mutant dogs

Previous studies reported that the *Shank3* mutant dogs exhibit impaired social interaction with humans (Ren et al., 2024; Tian et al., 2023). Given that affective information conveyed through touch makes important contributions to social behaviors (Gallace and Spence, 2010), we hypothesized that the observed social deficits in *Shank3* mutants might be linked to atypical AT responses. To test this possibility, we designed a three-phase, heterospecific human to dog AT paradigm (Fig. 1A; also see Supplementary Video 1 and Video 2). During habituation (phase 1), the adult dog freely explored the rectangular test arena for 10 min (2.7 x 1.8 x 0.9 m, see Methods for details). In phase 2 (simple presence), the experimenter stood in a corner of the test arena, looking straight ahead for 30 sec and did not interact with the test dog in any way. In phase 3 (hand stroke), the experimenter gently petted the test dog for two minutes.

**Fig. 1.**
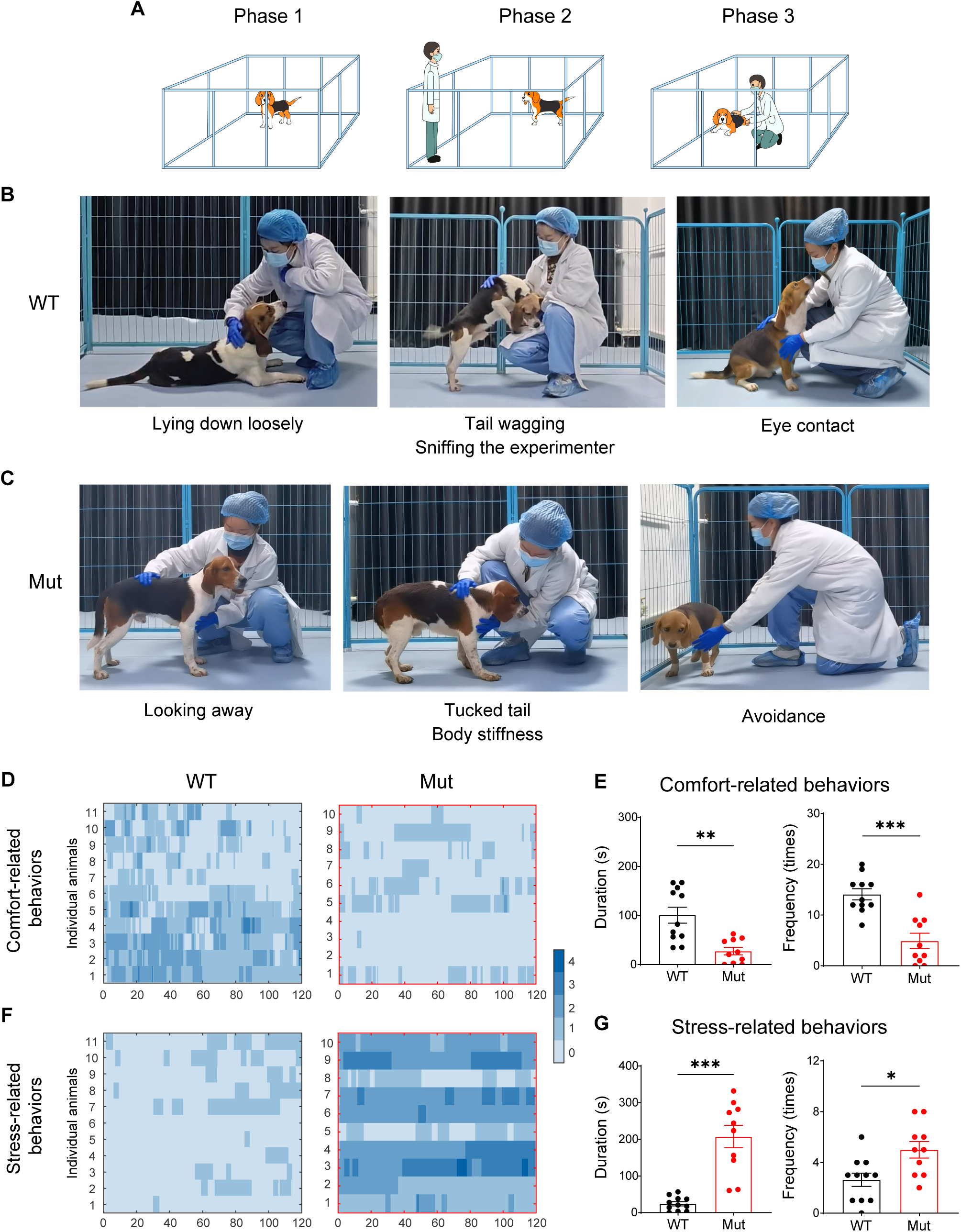
*Shank3* mutant dogs show aversion to AT, while WT dogs enjoy it. (A) An outline for the AT paradigm. Phase 1: habituation (10 min), phase 2: standing in the corner of the arena (30 s), phase 3: petting the dog (2 min). (B) The body language of pleasure in WT dogs during AT. Left: the WT dog lies down loosely. Middle: the WT dog wags its tail merrily and actively sniffs the experimenter. Right: the WT dog makes eye contact with the experimenter. The experimenter depicted in the images is one of the authors, and written informed consent was obtained for the publication of these photographs. (C) The body language of aversion in a *Shank3* mutant dog during AT. Left: the mutant dog exhibits eye avoidance and sustained fixation on distal environment. Middle: the mutant dog shows tucked tail and body stiffness with visible muscle tension. Right: the mutant dog exhibits avoidance behaviors withdrawal or flight. The experimenter depicted in the images is one of the authors, and written informed consent was obtained for the publication of these photographs. (D and F) Heat maps depicting the comfort-related behaviors (D) and stress-related behaviors (F) of individual dogs during 2-min phase 3 of AT paradigm. The different values in the scale bar represent the number of co-occurring comfort/stress-related behaviors. *n* = 11 for WT controls, and 10 for *Shank3* mutants. (E and G) *Shank3* mutants showed significantly altered duration and frequency of comfort-related (E) and stress-related (G) behaviors in phase 3 (hand stroke). Data are presented as mean ± SEM. \**p* < 0.05, \*\**p* < 0.01, \*\*\**p* < 0.001 by unpaired *t* test.

We quantified behaviors during phase 3 of the AT test, focusing on comfort-related behaviors and stress-related behaviors. Comfort-related behaviors included lying down loosely, tail wagging, sniffing the experimenter, and eye contact (Fig. 1B), while stress-related behaviors were looking away, tucked tail, body stiffness, and avoidance (Fig. 1C). *Shank3* mutant dogs enjoyed AT less, as indicated by remarkably reduced duration and frequency of comfort-related behaviors in phase 3 (Fig. 1D, 1E and Supplementary Table 1). Consistently, mutant dogs exhibited a higher level of stress to AT, as compared with WT dogs; the duration of stress-related behaviors was 207.70 ± 30.70 s, significantly longer than 24.39 ± 5.77 s of WT, as was the frequency of stress-related behaviors in *Shank3* mutants (5.00 ± 0.65 versus 2.64 ± 0.53 times/2 min) (Fig. 1F, 1G and Supplementary Table 2). Together, these results demonstrate that *Shank3* mutant dogs showed aversion to gentle stroke, while WT dogs enjoyed it. This finding is consistent with aversion to AT observed in ASD patients (Baranek, 1999; Mammen et al., 2015; Wiggins et al., 2009).

### The dmPFC encodes both affective touch and fast touch in WT dogs

To characterize the neuronal activity patterns in high-order cortical regions during AT, we developed a protocol using a 64-channel ultra-flexible neural probe (UFP) to record single-unit activities in the dorsal medial prefrontal cortex (dmPFC) (Fig. 2A–2C), a key region involved in emotion processing (Dixon et al., 2017; Etkin et al., 2011; Schirmer and Adolphs, 2017). The anatomical location and histological staining of the Beagle dmPFC were presented in Fig. S1A–C. The neural probe ensures minimal tissue damage and long-term signal stability via a minimized and flexible polyimide-gold sandwich construction as previously documented (Guan et al., 2019). As shown in Fig. S2A, single-unit waveforms from the same channel maintained highly stable over 26 days of recording (Pearson *r* > 0.80), both in WT and *Shank3* mutant dogs, over one year after electrode implantation. Beyond waveform stability, single-unit yield was also consistent across the 26-day recording period in both genotypes (Fig. S2B and S2C). Fig. 2D and 2E show the recording setup and the AT recording protocol, respectively. Dogs were petted with a stroking speed of 6–8 cm/s, previously shown to induce pleasant feelings in humans (Loken et al., 2009). This speed is also the speed with which one typically pets dogs. We then applied a faster stroking speed of 20–24 cm/s, referred to as fast touch (FT), which is perceived as less pleasant by humans (Loken et al., 2009). We then analyzed well-isolated spikes (Fig. 2C) aligned at the beginning of the stimuli when the dog received AT or FT from an experimenter. In the AT condition, approximately 18.38% of neurons in WT controls showed elevated firings while 17.65% of neurons showed the opposite (25 vs. 24 of 136 neurons; Fig. 2F), when compared with the resting-state firing rates. In the FT condition, the proportions for the two directions of responses were the same (16.18%, 22 of 136 neurons; Fig. 2I). These data indicate that the proportions of enhanced and suppressed neurons in the dmPFC of WT dogs during the two types of touch were similar.

**Fig. 2.**
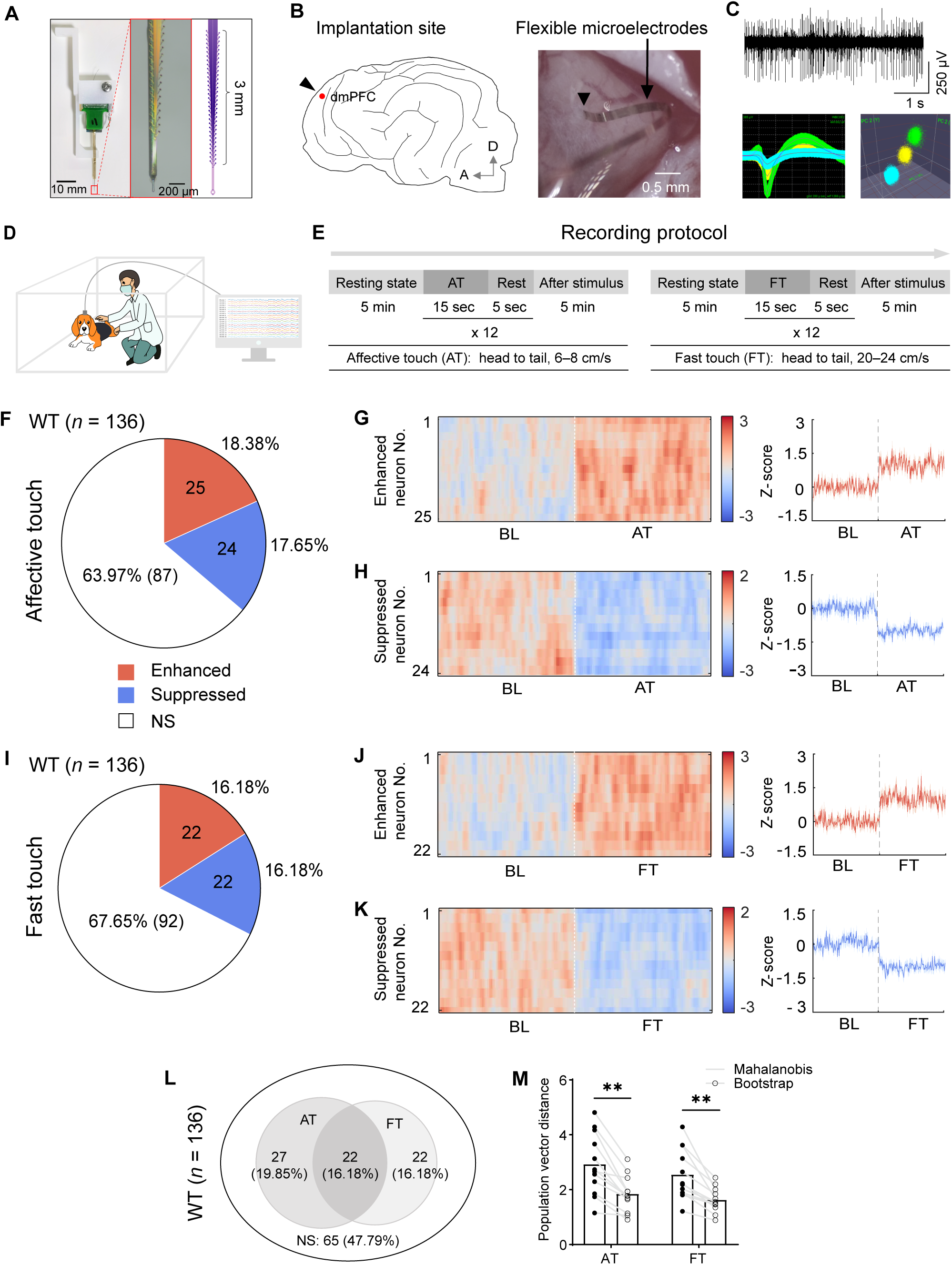
Neuronal responses to gentle tactile stimuli in the dmPFC of WT dogs. (A) Left: the ultra-flexible probe (UFP) for single-unit recordings; scale bar: 10 mm. Middle: zoom-in-view in the red box in the left; scale bar: 200 μm. Right: schematic diagram of the 3-mm-long linear contacts in the front of UFP. (B) Ultra-flexible electrode implantation in the dog brain. Left: schematic diagram illustrating the cortical location of implanted UFP into the dog dmPFC; A: anterior, D: dorsal. Right: image of open dura window showing the cortical surface after UFP implantation. Arrowheads: UFP insertion site; arrow: external connecting wires; scale bar: 0.5 mm. (C) Example of 250 Hz highpass continuous data of spike recordings in the dmPFC (top). Example of spike waveforms (bottom left) and the separation of single units in the principal component 3D space (bottom right). (D) Schematic diagram of single-unit recording setup. (E) The time line of single-unit recording protocol. (F, I) Proportion of neurons with enhanced, suppressed and non-significant (NS) response to AT (F) and FT (I) in the dmPFC of WT dogs. The significance was set at *p* < 0.05 by Wilcoxon rank sum test, two-sided, *n* = 136 neurons recorded. (G, H, J, K) Left: response profiles of neurons with enhanced and suppressed response to AT (G, H) and FT (J, K) aligned at the stimuli onset in the dmPFC of WT dogs. Right: response profiles calculated as z-scores in different conditions. (L) Venn diagram representing the overlap in coding properties between AT-coding neurons and FT-coding neurons in the dmPFC of WT dogs. (M) Population vector response between original and shuffled data using Mahalanobis distance for different tactile stimuli. The shuffling was performed using the bootstrap method. Two-way ANOVA followed by Bonferroni’s post hoc test. *n* = 12 sessions for both AT and FT. Data are presented as mean ± SEM. \*\**p* < 0.01.

We next normalized and averaged the firing rates of neurons that responded to each type of the stimuli. Both enhanced (Fig. 2G and 2J) and suppressed (Fig. 2H and 2K) firings were observed when the dog received the two types of tactile stimuli. We further found that 16.18% of all recorded neurons were responsive to both AT and FT in WT dogs (22 of 136 neurons; Fig. 2L). Among the subset of neurons that were responsive to either AT or FT, 30.99% (22 of 71) of neurons responded to both types of stimuli (Fig. 2L). To analyze the neural population responses to each type of stimuli, we measured the Mahalanobis distance (Durstewitz et al., 2010; Leutgeb et al., 2004), using population vectors composed of simultaneously recorded neurons. Population responses to AT and FT were significantly stronger than those from trial-shuffled bootstrap controls (stimulus and baseline trial labels randomly permuted across trials) (Fig. 2M), suggesting that both stimuli reliably evoked population-level neural responses beyond chance level. In summary, dmPFC neurons showed robust responses to AT and FT, supporting that dmPFC plays a key role in processing emotionally distinct tactile information.

### Loss of *Shank3* disrupts AT neuronal encoding and oscillatory activities in the dmPFC

In the dmPFC of *Shank3* mutant dogs, a smaller proportion of recorded neurons responded to AT (17.77% in mutants, as compared with 36.03% in WT dogs). Of these, only 4.61% (7 of 152) of neurons showed enhanced firing, while 13.16% (20 of 152) of neurons showed suppressed firing to AT, compared with the baseline level during resting state (Fig. 3A). In contrast, during FT, the proportion of responsive neurons was comparable between *Shank3* mutant and WT dogs (25.00% of 152 neurons versus 32.36% of 136 neurons; Fig. 3A and Fig. 2I). We further found that the proportion of neurons encoding both AT and FT in *Shank3* mutant dogs was lower than the WT controls (6.58% versus 16.18%; Fig. 3B and Fig. 2L).

**Fig. 3.**
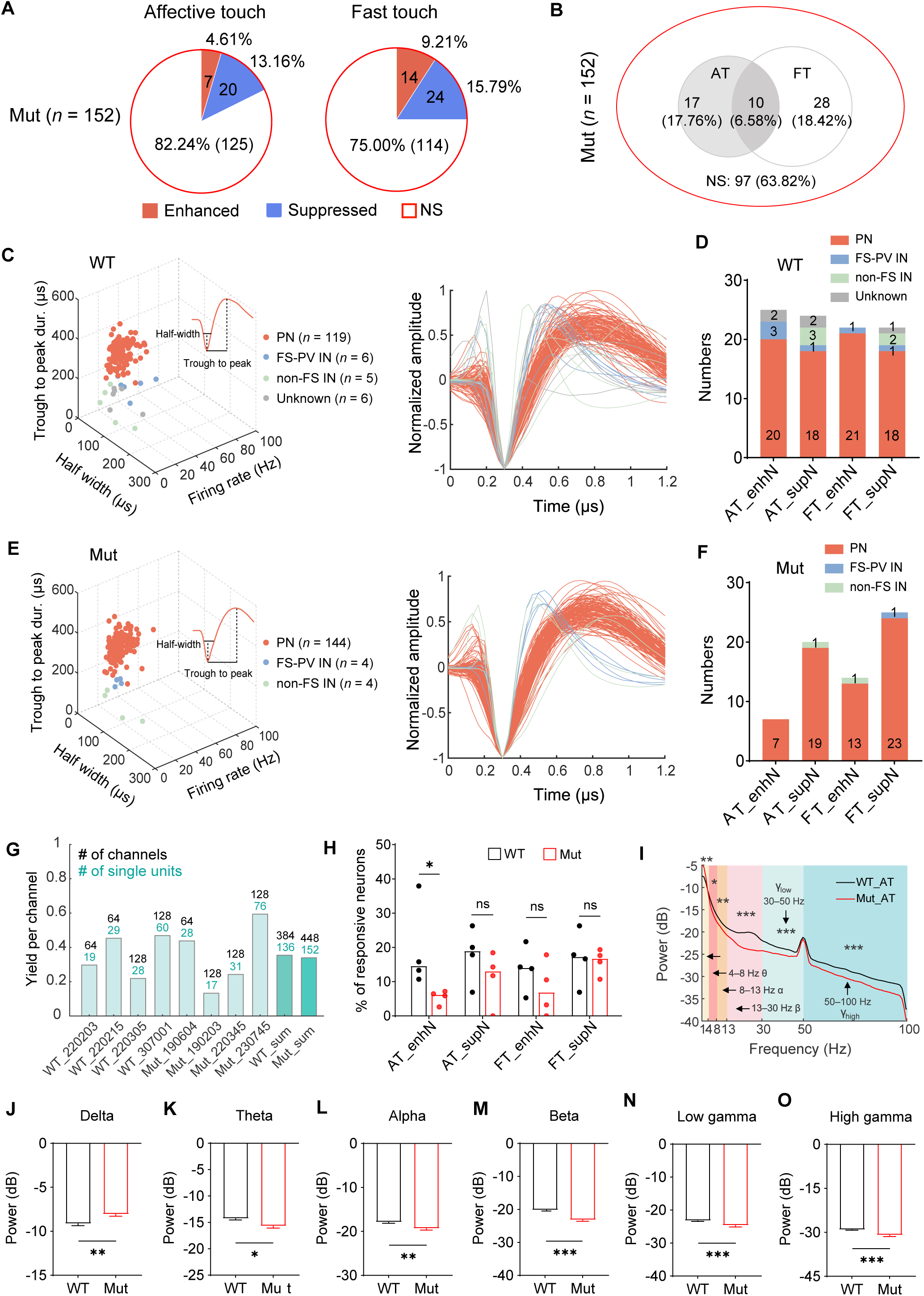
Impaired AT encoding in individual neurons and altered oscillatory activities in the dmPFC of *Shank3* mutant dogs. (A) Proportion of neurons with enhanced, suppressed and non-significant response AT (left) and FT (right) in the dmPFC of *Shank3* mutant dogs. The change of significance was set at *p* < 0.05 by Wilcoxon rank sum test, two-sided, *n* = 152 neurons. NS: non-significant. (B) Venn diagram representing the overlap in coding properties between AT-coding neurons and FT-coding neurons in the dmPFC of *Shank3* mutant dogs. (C) Left: all recorded dmPFC neurons in WT dogs were classified as PN, FS-PV IN, non-FS IN and unknown neurons; inset: average waveform of an example PN. Right: the average and normalized waveform of all recorded neurons in WT dogs. *n* = 136 neurons recorded. (E) The same as (C) but for *Shank3* mutant dogs. *n* = 152 neurons recorded. (D, F) Distribution of neuronal types encoding AT and FT in WT (D) and *Shank3* mutant dogs (F). AT_enhN: AT-responsive neuron with significant firing rate enhancement; AT_supN: AT-responsive neuron with significant firing rate suppression; FT_enhN: FT-responsive neuron with significant firing rate enhancement; FT_supN: FT-responsive neuron with significant firing rate suppression. (G) Single-unit recording yields in each dog. The numbers above each column indicate the number of channels and the number of well-isolated single units. The last two columns represent the summations from all WT and *Shank3* mutant dogs. (H) Ratio of neurons with enhanced or suppressed responses to AT or FT in WT and *Shank3* mutant dogs. (I) Group-level spectral powers of brain oscillations in the dmPFC of *Shank3* mutant (red, *n* = 4) and WT (black, *n* = 4) dogs during AT. Spectral powers at different frequency bands including delta: 1–4 Hz, theta: 4–8 Hz, alpha: 8–13 Hz, beta: 13–30 Hz, low gamma: 30–50 Hz, and high gamma: 50-100 Hz. (J–O) Comparisons of spectral powers at different frequency bands from the dmPFC of WT (black) and *Shank3* (red) mutant dogs during AT. Data are presented as mean ± SEM. \**p* < 0.05, \*\**p* < 0.01, \*\*\**p* < 0.001 by Mann-Whitney *U* test.

dmPFC contains both glutamatergic excitatory neurons and GABAergic inhibitory interneurons (Anastasiades and Carter, 2021). To further explore the cell-type specific responses to AT, we categorized touch-sensitive neurons based on spike features, into narrow-spiking (NS; trough to peak duration 239.62 ± 6.57 μs), putative inhibitory interneurons (IN), and wide-spiking (WS; trough to peak duration 474.50 ± 2.87 μs), putative pyramidal neurons (PN) (Fig. 3C and 3E) as previously described (Asadi et al., 2025; Dai et al., 2023; Hussar and Pasternak, 2009; Kim et al., 2016). The inhibitory INs were further classified into fast-spiking (average firing rate >10 Hz) parvalbumin (FS-PV) INs and non-FS INs based on neuronal firing rates (Fig. 3C and 3E) (Kim et al., 2016). The clustering analysis showed that the most cells recorded in WT and *Shank3* mutant dogs were excitatory pyramidal neurons. Among the 136 neurons recorded in WT dogs, there were 119 PNs, 6 FS-PV INs, 5 non-FS INs, and 6 undefined cells (Fig. 3C). In *Shank3* mutant dogs, of 152 recorded neurons, 144 were PNs, 4 were FS-PV INs, and 4 were non-FS INs (Fig. 3E). These results indicated that PNs in dmPFC may contribute substantially to AT processing.

We then analyzed the cell type composition of neurons that encoded AT and FT and found that AT-encoding neurons (enhanced or suppressed responses to AT) in the dmPFC of WT dogs comprised a small subset of FS-PV INs (4 out of 49 neurons; Fig. 3D). In contrast, no FS-PV INs (0 out of 27 neurons) were involved in AT processing in *Shank3* mutant dogs (Fig. 3F). To examine whether sampling bias could contribute to this difference in cell type composition, we quantified the single-unit yield for each dog. We identified 136 and 152 single units from 384 and 448 channels in WT and mutant dogs, respectively, corresponding to comparable yields of 0.35 and 0.34 single units per channel (Fig. 3G). In addition, the proportion of neurons with enhanced responses to AT in *Shank3* mutant dogs was significantly reduced compared with WT dogs (6.17% versus to 14.56%; Fig. 3H). In contrast, no significant difference was observed in the proportion of neurons with suppressed responses to AT between the two genotypes. Similarly, there were no significant differences in the proportions of neurons responsive to FT between WT and mutants (Fig. 3H). In summary, *Shank3* mutant dogs exhibited a selective reduction in the proportion of neurons with enhanced responses to AT in the dmPFC.

Brain oscillations reflect spontaneous neural activity and are characterized by spectra at different frequency bands, each has been associated with different cognitive processes (Han et al., 2021; Ward, 2003; Xing et al., 2012). To further investigate the neural responses to AT, we compared AT-induced oscillatory activities between WT and *Shank3* mutant dogs. Spectral analysis revealed distinct differences in the dmPFC of the two groups, as shown in Fig. 3I. Specifically, *Shank3* mutant dogs exhibited significantly lower spectral power during AT across the theta, alpha, beta, low-gamma, and high-gamma frequency bands (Fig. 3K–3O). In contrast, delta activity was markedly higher in *Shank3* mutant dogs compared with WT controls (Fig. 3J). Together, these findings show impaired neuronal encoding and altered oscillatory dynamics during AT in the dmPFC of *Shank3* mutant dogs.

### PTZ rescues altered behavioral and neural responses to AT in *Shank3* mutant dogs

We recently showed that GABA_A_ receptor antagonist pentylenetetrazole (PTZ) significantly improves tactile threshold and promotes social interaction in *Shank3* mutant dogs (Shi et al., 2025b). To test whether PTZ could ameliorate AT aversion in *Shank3* mutant dogs, we conducted a two-phase behavioral rescue experiment (Fig. 4A). In Phase 1 (4 days total), test dogs received intramuscular saline on days 1 and 2 without AT. On day 3, test dogs received saline injection, followed by PTZ administration on day 4, with AT test conducted 30 min post-injection in both conditions. To control for potential habituation effects from repeated AT tests, we performed a phase 2 experiment. After a 7-day washout period, saline was administrated on days 11–14, with AT test performed 30 min post-injection only on days 13 and 14. We administrated PTZ intramuscularly to *Shank3* mutants at 1.5 mg/kg; this dose is an order of magnitude lower than the reported seizure-inducing dose, and we did not observe any signs of seizure or other side effects in the previous studies (Shi et al., 2025a; Shi et al., 2025b) and the present study. Behavioral analysis revealed that PTZ treatment did not increase the duration of comfort-related behaviors in *Shank3* mutant dogs during AT (Fig. 4B). However, the duration of stress-related behaviors in *Shank3* mutant dogs during AT was significantly ameliorated by PTZ treatment (day 4 versus day 3) in phase 1, while the saline control in phase 2 showed no rescue effect (day 14 versus day 13) (Fig. 4C). In addition, PTZ did not significantly affect the frequency of comfort-related and stress-related behaviors in *Shank3* mutants (Fig. 4D and 4E).

**Fig. 4.**
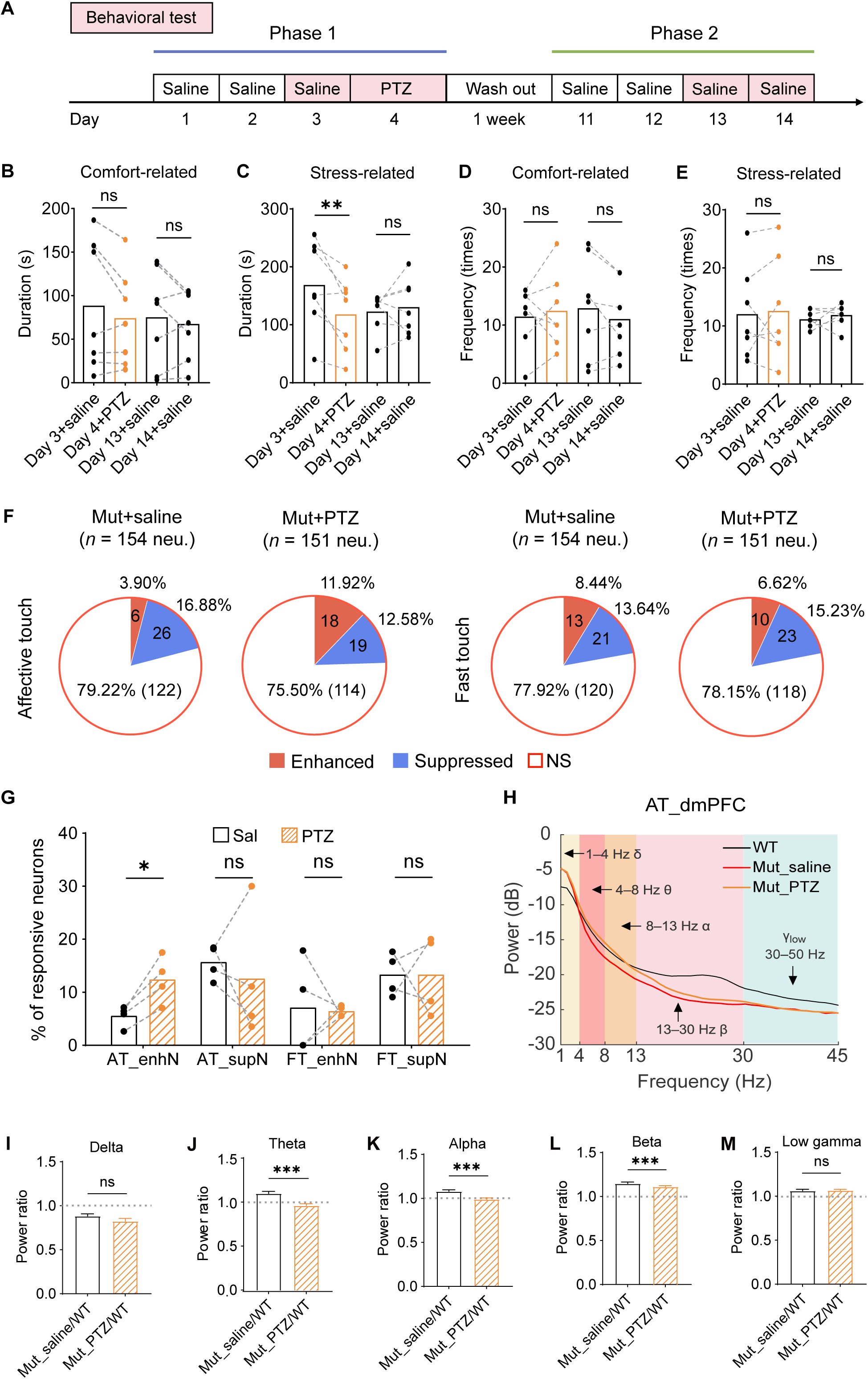
PTZ rescues the AT deficits in *Shank3* mutant dogs. (A) Timeline of drug treatment. The test dogs receive saline (0.9% NaCl) and no AT behavioral test on days 1, 2, 11 and 12. On days 3, 4, 13 and 14, the test dogs perform AT behavioral test at 30 min after saline or PTZ administration. (B) *Shank3* mutant dogs showed no significant changes in the duration of comfort-related behaviors during AT after PTZ administration compared with the saline group. *n* = 7. (C) The duration of stress-related behaviors in *Shank3* mutant dogs during AT was significantly reduced after PTZ administration compared with the saline group. *n* = 7. (D, E) *Shank3* mutant dogs showed no significant changes in the frequency of comfort-related behaviors (D) and stress-related behaviors (E) during AT after PTZ administration compared with the saline group. *n* = 7. (F) Proportion of neurons with enhanced, suppressed and non-significant response to AT (left panel) and FT (right panel) in the dmPFC of *Shank3* mutant dogs after saline and PTZ treatment. *n* = 154 and 151 neurons in the saline and PTZ groups, respectively. (G) Ratio of neurons with enhanced or suppressed response to AT or FT in *Shank3* mutant dogs following saline or PTZ administration. *n* = 4. (H) Group-level spectral powers of brain oscillations in the dmPFC of WT (black, *n* = 4) and *Shank3* mutant (saline: red, PTZ: yellow, *n* = 4) dogs during AT. Spectral powers at different frequency bands including delta: 1–4 Hz, theta: 4– 8 Hz, alpha: 8–13 Hz, beta: 13–30 Hz, and low gamma: 30–45 Hz. (I–M) Comparisons of power ratios between *Shank3* mutants treated with saline or PTZ and WT controls at different frequency bands in the dmPFC during AT. Data are presented as mean ± SEM. \**p* < 0.05, \*\**p* < 0.01, \*\*\**p* < 0.001, ns: no significance, by two-way ANOVA followed by Bonferroni’s post hoc test.

GABA_A_ receptor antagonists have been reported to inhibit the action of GABA and consequently increase the excitability of PNs (Krogsgaard-Larsen et al., 1994; Sigel and Steinmann, 2012). To investigate whether PTZ could rescue the disrupted neuronal encoding of AT in *Shank3* mutant dogs, we performed single-unit recording before and after PTZ treatment. PTZ treatment increased the proportion of dmPFC neurons with enhanced responses to AT in *Shank3* mutants (3.90% for saline and 11.92% for PTZ; Fig. 4F). In contrast, the proportion of neurons with suppressed responses to AT was comparable between groups (16.88% for saline and 12.58% for PTZ; Fig. 4F). Consistent with this pooled results, per-animal statistics confirmed that PTZ significantly increased the proportion of neurons with enhanced responses to AT, while the proportion of AT-suppressed neurons was not significantly altered (Fig. 4G). For FT, PTZ did not significantly affect the proportions of neurons responding to FT compared with saline treatment in the dmPFC of *Shank3* mutants (Fig. 4F and 4G). Furthermore, PTZ administration restored the reduced spectral powers of theta, alpha, and beta but not delta and low gamma bands in *Shank3* mutant dogs, compared with saline treatment control (Fig. 4H–4M). Collectively, the rescue effects of the GABA_A_ receptor antagonist PTZ on the behavior, neuronal encoding and oscillations in *Shank3* mutant dogs during AT highlights the importance of GABAergic signaling in tactile processing and suggests that GABAergic signaling pathway might serve as a potential therapeutic target for correcting abnormal AT processing in patients with *SHANK3* mutations.

## Discussion

Taking advantage of the *Shank3* mutant dogs, which display aversion to AT as observed in ASD patients (Baranek, 1999; Mammen et al., 2015; Wiggins et al., 2009), we established a long-term single-unit recording system to explore the neural basis of AT. We showed that loss of *Shank3* disrupts AT-evoked brain oscillations and neural encoding in individual neurons in the dmPFC, especially the excitatory pyramidal neurons. Importantly, the AT aversion and neural encoding phenotypes in *Shank3* mutant dogs can be rescued by the GABA_A_R antagonist PTZ. Together, these results reveal the neural mechanisms underlying AT deficits associated with *Shank3* mutations and provide a foundation for developing targeted interventions for tactile processing impairments in ASD.

A major obstacle in dissecting the neural basis of AT is the lack of a robust, quantifiable experimental paradigm. Here, we introduce a cross-species human to dog AT paradigm that bridges this gap: it is based on a strong, naturalistic bond between dogs and humans, enabling efficient social interactions. In this paradigm, the duration, speed, and force of social stroke are tightly controlled in a quantitative manner. Furthermore, touch sensation in dogs bears closer resemblance to that in humans, as skin is the primary target for tactile stimulation (Hashimoto et al., 1986). These features, combined with dogs’ evolved sensitivity to human social cues (Hare and Tomasello, 2005), establish dogs as a powerful cross-species model for understanding AT from sensory input to emotional valence.

Previous studies demonstrated that the PFC is implicated in processing pleasant touch in humans (Francis et al., 1999; Gordon et al., 2013), consistent with its well-established role in high-order cognitive and social-emotional functions. Through single-unit recording, we identified a group of cells in WT dogs’ dmPFC that respond strongly to AT, suggesting that dmPFC encodes affective tactile information. The relative contribution of distinct neuronal classes to sensory stimuli in the PFC vary across species. For example, in mice performing a sensory discrimination task, PV+ interneurons are the only class with robust responses to sensory stimuli, whereas pyramidal neurons display greater functional heterogeneity with varied response profiles in the PFC (Pinto and Dan, 2015). In the present study, we observed significantly fewer AT-enhanced neurons in the dmPFC of *Shank3* mutant dogs, particularly the excitatory pyramidal neurons. This observation is consistent with SHANK3 being an abundant synaptic scaffolding protein primarily located in the postsynaptic density (PSD) of excitatory glutamatergic synapses (Jiang and Ehlers, 2013; Monteiro and Feng, 2017; Yang et al., 2024). This cell-type-specific deficit in AT encoding likely impairs local computation within the dmPFC microcircuit, thereby compromising the regulation of pleasure-related behaviors. Our findings in dog models are consistent with a diminished response to pleasant stimuli in the PFC of ASD patients by fMRI (Kaiser et al., 2016). Cortical hypo-reactivity to AT may contribute to diminished reward perception from social touch, which in turn contributes to social interaction deficits in ASD.

The rescue of frequency-specific oscillatory deficits in the dmPFC by PTZ provides crucial insights into the neural basis of AT processing. The mPFC is a key node in networks affecting affective evaluation, cognitive control, and reward processing (Chen et al., 2021; Rudebeck et al., 2013). Among the rescued frequency bands, theta oscillation in the dmPFC is strongly implicated in emotional valence processing, cross-regional communication during fear learning, and the integration of affective information (Chen et al., 2021). Alpha and beta rhythms in prefrontal regions are associated with internal state regulation, top-down sensory prediction, and the encoding of reward value (Helfrich and Knight, 2016; Klimesch, 2012). The restoration of these low-frequency rhythms (theta, alpha and beta) in *Shank3* mutants suggests that PTZ ameliorates the deficit in assigning affective salience and generating the pleasant, prosocial sensation intrinsic to AT. This effect likely stems from PTZ’s modulation of network excitability within the mPFC and its connected limbic-striatal circuits that subserve the cognitive and emotional appraisal of sensory events (Christakou et al., 2004; de Kloet et al., 2021).

In contrast, PTZ failed to rectify the oscillatory anomalies in the delta and gamma bands. Delta oscillations (1–4 Hz) were elevated in *Shank3* mutants and remained unaltered after PTZ treatment. Recent evidence indicates that optogenetic inhibition of cortical PV+ interneurons enhances delta power, whereas their activation attenuates it, suggesting that PV+ interneuron function contributes to the maintenance of normal delta-band dynamics (Hu et al., 2025). High-frequency gamma oscillations (30–100 Hz) are fundamental for the precise temporal encoding of low-level sensory features, such as pain intensity (Yue et al., 2025). The reduced gamma oscillations suggests a fundamental deficit in the cortical circuitry responsible for processing the physical properties of the tactile stimulus itself (e.g., velocity, intensity, frequency) (Bessaih et al., 2018). It is worth noting that the frequency-specific rescue effect may align with the distinct cellular origins of different oscillations. Low-frequency rhythms are more closely linked to excitatory pyramidal neuron synchronization (Buzsaki and Draguhn, 2004; Einevoll et al., 2013). In contrast, the generation of gamma rhythms critically requires INs, especially the PV+ INs (Buzsáki and Wang, 2012). Therefore, the persistent gamma deficit in *Shank3* mutants may reflect PTZ’s inability to restore the interneuron-dependent gamma rhythmogenesis. Similarly, the unaltered delta power may be related to PTZ’s limited effect on the interneuron networks that regulate delta dynamics. In summary, the rescue effects of PTZ likely arise from its regulation of pyramidal neuron-involved circuits, selectively restoring theta, alpha, and beta oscillations essential for affective integration in the dmPFC.

There are a few limitations in the current study. For one, we only investigated cortical processing of AT. Future work needs to explore the role of subcortical nuclei, such as nucleus accumbens (NAc) and ventral tegmental area (VTA), which have long-range connections with the PFC and are involved in reward processing. A second limitation lies in the fact that our investigation focused on adult dogs. Given the importance of AT in social development, future research would should explore the role of early-life experiences, particularly during critical social development windows, in correcting abnormal AT processing in *Shank3* mutant dogs. Though the causality of decreased gamma oscillation during AT in *Shank3* mutants remains to be further investigated, our current data demonstrate that it plays a critical role in AT processing. Based on this and other studies (Li et al., 2025; Tian et al., 2023; Yuan et al., 2025), *Shank3* mutant dogs faithfully recapitulate a subset of the clinical features of ASD and allow us to further dissect a mechanistic link between impaired AT processing and social impairments in ASD.

## Acknowledgments

We thank Jianhong Luo from Zhejiang University for the discussion and Chengyu Li’s team from Lingang Laboratory, Shanghai, China, for their assistance in brain surgery of large animals; Mengcheng Liu from the National Center for Nanoscience and Technology, for technical help on single-unit recording; Yang Zhan from the Brain Cognition and Brain Disease Institute, Shenzhen Institute of Advanced Technology, Chinese Academy of Sciences for the population vector analysis of spike data. This work was supported by the National Natural Science Foundation of China (Grant No.32394030) and Wuhan Municipal Science and Technology Bureau (Grant No. 2024020702030125).

## Author contributions

Yanhe Zhou, Minna Dan, Yue He, Xiang Yu, Jinfen Wang, Dajun Xing and Yong Q. Zhang designed the experimental paradigms. Yanhe Zhou, Minna Dan, and Yue He performed experiments. Yanhe Zhou, Minna Dan, and Lintao Jia developed surgical procedure for the single-unit recording technique in dogs. Yanhe Zhou, Minna Dan and Yue He analyzed data. Ying Fang and Jinfen Wang designed and provided the ultra-flexible neural probes. Yanhe Zhou wrote the manuscript. Kun Guo, Li Hu, Xiang Yu, Dajun Xing, and Yong Q. Zhang revised the manuscript.

## Data availability statement

The data that support the findings of this study are available from the corresponding author upon reasonable request.

## Materials and Methods

### Animal husbandry

All adult male Beagle dogs, both *Shank3* mutant and WT, weighing between 10 and 16 kg, were housed individually in home cages (1 x 1 x 1 m^3^) under temperature and humidity controlled conditions (22–24°C, 40–60% humidity). All dogs received food twice a day and water ad libitum and were kept in a 12-hour day-night cycle (lights on from 07:00 to 19:00). The life experience of each animal was kept as similar as possible to minimize individual variability. All surgical and experimental procedures, as well as animal care and handling, were approved by the Animal Ethics and Welfare Committee of Hubei University (20230066) and of the Institute of Genetics and Developmental Biology, Chinese Academy of Sciences (AP2024025).

### Human affective touch on dogs

The test of AT was conducted in a puppy pen of 2.7 x 1.8 x 0.9 m (length x width x height). Test dogs were individually habituated to the puppy pen for 10 min in the absence of an experimenter (phase 1). Each dog was then allowed to interact freely with an experimenter standing in a corner of the puppy pen for 30 s (phase 2). The experimenter then squatted and stroked the test dog for 2 min (phase 3). The experimenter (one of the authors) was unfamiliar with all the test dogs and completed the whole AT assay. All behaviors were recorded on video throughout the test.

We quantified the dog’s behaviors only in phase 3. The main behaviors we focused on are comfort-related and stress-related behaviors. Comfort-related behaviors include lying down loosely (sprawling out on the floor with its body visibly at ease and muscles relaxed, or resting in a casual pose with its belly exposed and limbs stretched in different directions), tail wagging, sniffing the experimenter, and eye contact (looking at the experimenter’s face) (Fig. 1B). Stress-related behaviors were looking away (turning its head aside and focusing intently on something else in the room), tucked tail, body stiffness (standing unmoving with its body rigid and tense, muscles visibly tight), and avoidance (retreating slowly or moving away evasively) (Fig. 1C).

### Ultra-flexible neural probe designed for dogs

A 64-channel ultra-flexible neural probe (UFP) was made using a standard micro-fabrication process (Guan et al., 2019). The probe consists of a proximal bonding pad section for electrical connection to the neural signal recording system and a distal deep implantation section for recording neural activity. It features a sandwich structure with two polyimide insulating layers and a gold conductive layer, exposing bonding pads and recording sites. The implantation section has a tapered design, transitioning from a fine tip to a wider base to minimize tissue damage. The impedance of the recording sites is 50–100 kΩ at 1 kHz in saline solution. These sites are linearly distributed along a 3-mm depth, with a 90-µm spacing between adjacent sites. Each recording site has a circular shape with a diameter of 20 μm. An auxiliary implantation hole is located at the distal end of the implantation section. To minimize implantation damage, a small-footprint tungsten wire was assembled with the implantation hole to ensure precise implantation. After the implantation, the tungsten wire was withdrawn from the brain and the probe remained in the targeted brain region.

### Surgical procedures

Surgery for electrode implantation was performed under general anesthesia and strictly sterile conditions as previously described (Wu et al., 2023). The test dog was anesthetized by simultaneous administration of dexmedetomidine hydrochloride (5 μg/kg, intramuscular), Zoletil 50 (1 mg/kg, intramuscular), and propofol (5 mg/kg, intravenous). The anesthetized dogs were maintained by endotracheal intubation with 3% isoflurane, and mounted on a stereotaxic apparatus. Electrocardiography (ECG) and oxygen saturation monitors were attached to the animals to monitor vital signs. After the scalp sterilization procedures, a scalp incision of 6 cm was made at the prefrontal area, exposing the skull beneath. The cranial windows of 10 × 10 mm and 4 × 4 mm were created on the first and second layer of skull using a dental drill, respectively. The dura mater was then incised to expose the underlying cortex.

The UFPs were implanted one by one in a modularized manner. A UFP was first attached to a customized 3D printed resin holder mounted on a micromanipulator arm, then slowly propelled into the target brain area, with an implant depth of about 3.5 mm. This insertion depth was chosen based on the average thickness of the cortex and MRI data of the test dog. Following the insertion of the shuttle wire, all the recording sites were meticulously examined under a microscope to confirm their successful implantation into the cortex, with the proximal contact just beneath the cortical surface. The tungsten shuttle wire was then retracted manually. After removing the shuttle wire, all electrode sites were examined again under a microscope to verify that they all remained in place and were not pulled out of brain tissue. Then, the first UFP was detached from the manipulator to load the second UFP onto micromanipulator. The same procedure was repeated until the second UFP was implanted. Each dog received either one or two UFPs containing 64 or 128 channels in total (Fig. 3G), with implantation sites 1–2 mm apart when two probes were used.

After implantation, the cranial window was sealed with the human fibrin sealant (FIBINGLUEAAS, Shanghai, China) and Kwik-Sil (World Precision Instruments, Sarasota, FL). Six to eight skull screws were placed in the drilled craniotomies, on which the customized chamber was fixed using light-curing dental acrylic (3M). A subsequent layer of dental acrylic was applied over the craniotomy and surrounding skull surface to firmly fix the silicon carrier chips to the skull and chamber. Finally, a customized cap was placed to cover the chamber. The dog with implanted UFPs was given postsurgical intramuscular injection of antibiotics every day for one week.

### AT and FT stimuli for *in vivo* electrophysiology

We used a stroking speed of 6–8 cm/s (i.e., AT), previously shown to be most effective at activating low-threshold mechanosensitive C-tactile fibers that transduce sensory information from the periphery, to induce pleasant feelings in humans (Loken et al., 2009). This speed is also the normal speed when we usually pet the dogs. For control, we used a faster stroking speed of 20–24 cm/s (i.e., FT) which is known to induce a less pleasant perception in humans (Loken et al., 2009). Gentle stroking was performed by a trained experimenter who was blind to the genotype. The dog was gently stroked by the experimenter’s hand, moving from the neck to the lumbar region (about 24 cm stroking distance) at a specific speed (6–8 cm/s or 20–24 cm/s) and force (0.5–1 N). Force was assessed by pressing the hand against a balance. Each session included 5 min of resting state, 12 trials of AT or FT (15 s stroking interspersed with 5 s rest), and 5 min of post-trial rest (Fig. 2E). Six sessions were carried out per dog, comprising three AT sessions and three FT sessions, and the order of AT and FT was randomized across sessions to eliminate bias. All sessions were conducted between 9:00–11:00 am. In total, we performed 48 sessions on 4 WT and 4 *Shank3* mutant dogs for both AT and FT conditions.

### Neural recordings and single-unit analysis

Neural recordings were initiated following a 2-week recovery from the surgery. Spike activity was recorded in animals with restricted moving in square fences (0.9 m × 0.9 m) using Zeus data acquisition systems (Zeus, Nanjing, China). Recording was performed at a sampling rate of 30 kHz. A 250 Hz high-pass filter was applied to single-unit recordings. Spikes were identified when a minimum waveform reached an amplitude threshold of 3 standard deviations higher than the noise amplitude. Spike sorting was performed offline using Offline Sorter 4.7.1 (Plexon systems). Principal component scores were used to cluster units and the L-ratio and isolation distance were calculated. A group of waveforms was considered to be generated from one single unit if it was distinct from other clusters. Manual checking was then performed to ensure that similar spike waveforms were clustered and that the cluster boundaries were clearly separated. Clusters whose fraction of spikes with inter-spike intervals within 1 ms was higher than 1% of the total firing events were discarded. Only well-isolated units were included in the data analysis.

Single-unit spike data were converted to point processes at 1 ms with 0/1, and the firing rate was calculated using a 100 ms bin. Neurons with a mean firing rate of less than 0.5 Hz were not used. To evaluate the response of a neuron, a non-parametric rank sum test was used to compare the firing rates of baseline (resting state) and tactile stimuli periods across all trials. A *p* value less than 0.05 was used as the criterion for being responsive. To examine the response profiles for each stimulus, firing rate changes within each stimulus were averaged across trials. The firing rate was then converted to a z-score by normalizing the baseline.

To evaluate the stability of neural recordings over time, representative single units were selected from implanted UFPs in both WT (e.g., channel 52 from WT307001,) and *Shank3* mutant (e.g., channel 72 from Mut220345) dogs. The mean spike waveforms were extracted on each recorded day (spanning 26 days at least one year after implantation). Stable recording was inferred from minimal day-to-day variations in waveform morphology relative to the initial recording, as quantified by Pearson correlation coefficients (*r,* with *r* = 1 set as baseline). We also quantified the single-unit yield per channel to evaluate the efficiency of electrode recordings in capturing neuronal signals. The overall yield for each dog was defined as the total number of good units recorded divided by the total number of channels.

### Local field potential data analysis

The local field potential (LFP) signals were resampled to 1000 Hz and band-pass filtered between 1 and 100 Hz to remove low-frequency fluctuations and high-frequency noises. LFP data were analyzed in MATLAB version 2019a (Mathworks Inc., Natick, MA), along with the EEGLAB. The bad channels were excluded by visual inspection. Epochs were extracted using a window analysis time of 1 s during tactile stimulus (1.5–13.5 s after stimulus onset) and were baseline-corrected by the pre-stimulus 5-min interval. Epochs whose amplitude exceeded 8 standard deviations at any point of the time course were identified as contaminated by gross artifacts and were automatically discarded without further analysis. Less than 10% of the total number of epochs on average were removed from each dog, and the remaining epochs were averaged across trials. Power spectral density (PSD) describes how the power of a LFP is distributed over frequency. In analyzing the frequency content of the signal, the function PWELCH in MATLAB was performed on each epochs to estimate PSD via Welch’s method. The normalized PSD is defined as the PSD during tactile stimulation subtracting the PSD during the 5-min rest before stimulus across all trials. Frequencies from 1 to 100 Hz were divided into six bands: delta (1–4 Hz), theta (4–8 Hz), alpha (8–13 Hz), beta (13–30 Hz), low gamma (30–50 Hz) and high gamma (50–100 Hz).

### Population vector analysis

To calculate the population response, the population vector was calculated from the firing rates of all cells in the session. Only sessions in which at least eight cells were recorded simultaneously were used. We calculated the Mahalanobis distance between baseline (resting state) and during-stimulus (15 s after the onset of tactile stimulation). The calculation formula was as follows: 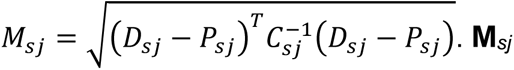 was the Mahalanobis distance between baseline and during stimulus population vectors in the j-th trial of stimulus AT and FT. **D***_sj_* was the during-stimulus population vector. **P***_sj_* was the baseline population vector. **C***_sj_* was the covariance matrix. We used the maximum Mahalanobis distance of all trials for each stimulus in each session.

To evaluate the responsiveness of the population vector, we used a bootstrap method to shuffle the trials containing the population vectors across the stimuli. The procedure was repeated for 1000 times in the same recording session and a shuffled Mahalanobis distance was calculated.

### Unit classification

The well-isolated units were first classified into wide-spiking (WS) putative pyramidal neurons and narrow-spiking (NS) INs using unsupervised cluster algorithm based on k-means method. The analysis was based on the three-dimensional space defined by each neuron’s half-spike width (trough to peak duration), half valley width and the mean firing rate at baseline. Spikes with shorter half-spike width, half valley width and higher firing rate were classified to be putative INs. The NS population was further classified into putative FS-PV INs (> 10 Hz) and non-FS INs based on baseline firing rate (Courtin et al., 2014; Kim et al., 2016).

### Pentylenetetrazole (PTZ) administration

PTZ, a GABA_A_ receptor antagonist, increases the duration of the closed state of the GABAA receptor by inhibiting the GABA-activated Cl⁻ current in a concentration-dependent manner, thereby maintaining neuronal excitability (Huang et al., 2001). To investigate whether the GABA_A_ receptor antagonist could rescue the aversive behaviors to AT in *Shank3* mutants, we administered PTZ intramuscularly at a concentration of 1.5 mg/kg, a dose that did not induce seizures, as documented in a previous study (Shi et al., 2025b). No other adverse effects were observed in dogs after PTZ administration. After saline or PTZ administration, the dogs were returned to their home cages for 30 min before performing experimental assays, which included AT behavioral assessment and the collection of electrophysiological data.

### Immunohistochemistry and image acquisition

Dogs were deeply anesthetized with 1 mg/kg Zoletil 50. Carotid artery perfusion was performed with PBS followed by 4% PFA. Coronal brain sections were cut with a freezing microtome (Leica CM 1950) at 30 μm. Sections were blocked in PBS containing 5% bovine serum albumin and 0.3% Triton X-100 for 1 h at 25°C, followed by incubation with primary antibody overnight at 4°C. After ample washes, sections were incubated with secondary antibodies for 2 h at 25°C. Neurons were identified using mouse anti-NeuN antibody (1:1000, Abcam, Cat# ab104224, RRID:AB_10711040). GFAP expression was identified using rabbit anti-GFAP antibody (1:400, Zhongshan Golden Bridge, Cat# ZA-0529). The following secondary antibodies (all from Thermo Fisher Scientific) were used at 1:500: donkey anti-Mouse Alexa Fluor 488 (Cat# A-21206, RRID:AB_2535792) and donkey anti-Rabbit Alexa Fluor 568 (Cat# A10042, RRID:AB_2534017). DAPI (1:3000, Thermo Fisher Scientific, Cat# D1306, RRID:AB_2629482) was used to stain nuclei. Images of immunostaining were acquired using PerkinEImer Vectra Polaris System with a 20x objective (N.A. = 0.75).

### Statistical analysis

Data are presented as mean ± standard error of the mean (SEM). The statistics were performed using Graphpad Prism 10. Unless otherwise specified, comparisons between two groups were tested using the unpaired two-sample Student’s t test. Multiple comparisons were carried out by one-way ANOVA (the Newman-Keuls test) or two-way ANOVA. In all figures: \**p* < 0.05, \*\**p* < 0.01, and \*\*\**p* < 0.001.

---

**Supplementary Fig. 1.**
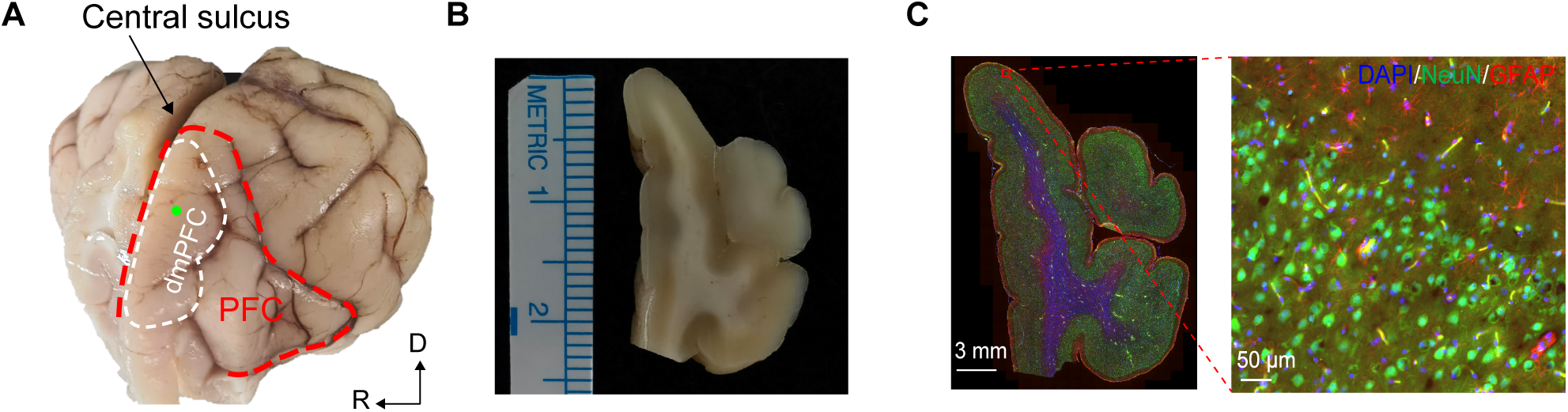
*In vivo* single-unit recording in the dmPFC of dogs. (A) The physical picture of part of the Beagle’s brain. The red dashed line demarcates the prefrontal cortex. The white dashed line indicates dmPFC. Green dot denotes UFP insertion site. R: right, D: dorsal. (B) Coronal section of the right hemisphere of the brain near the electrode implantation site, showing clear gray-white matter demarcation. The length of a small grid on the left ruler represents 1 mm. (C) Left: Immunofluorescence staining of neuronal (NeuN, green), astrocyte (GFAP, red), and nuclear DAPI (4’,6-diamidino-2-phenylindole, blue) markers in the coronal section of the right prefrontal lobe of a Beagle dog. Scale bar: 3 mm. Right: higher-magnification view of the boxed region in the left panel, revealing the cellular composition. Scale bar: 50 μm.

**Supplementary Fig. 2.**
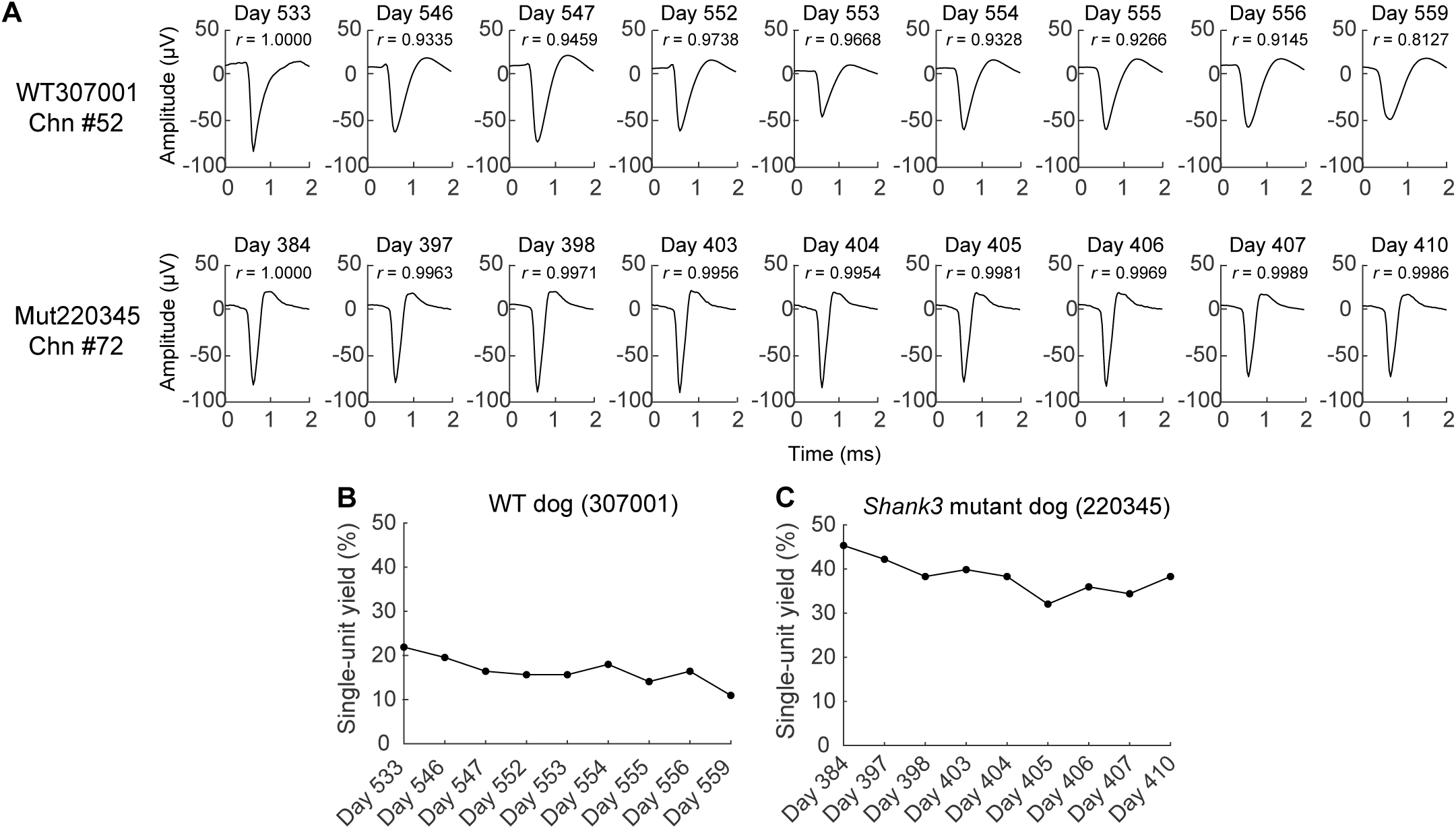
Stable single-unit recording in dogs at least one year after electrodes implantation. (A) Average waveforms of two representative units in WT (top) and *Shank3* mutant (bottom) dogs at least one year after the electrode array implantation. Each waveform represents the average of all spikes. (B, C) Single-unit yield (%) for the WT dog (307001) and the *Shank3* mutant dog (220345) at least one year after electrode array implantation.

**Supplementary Table 1.**
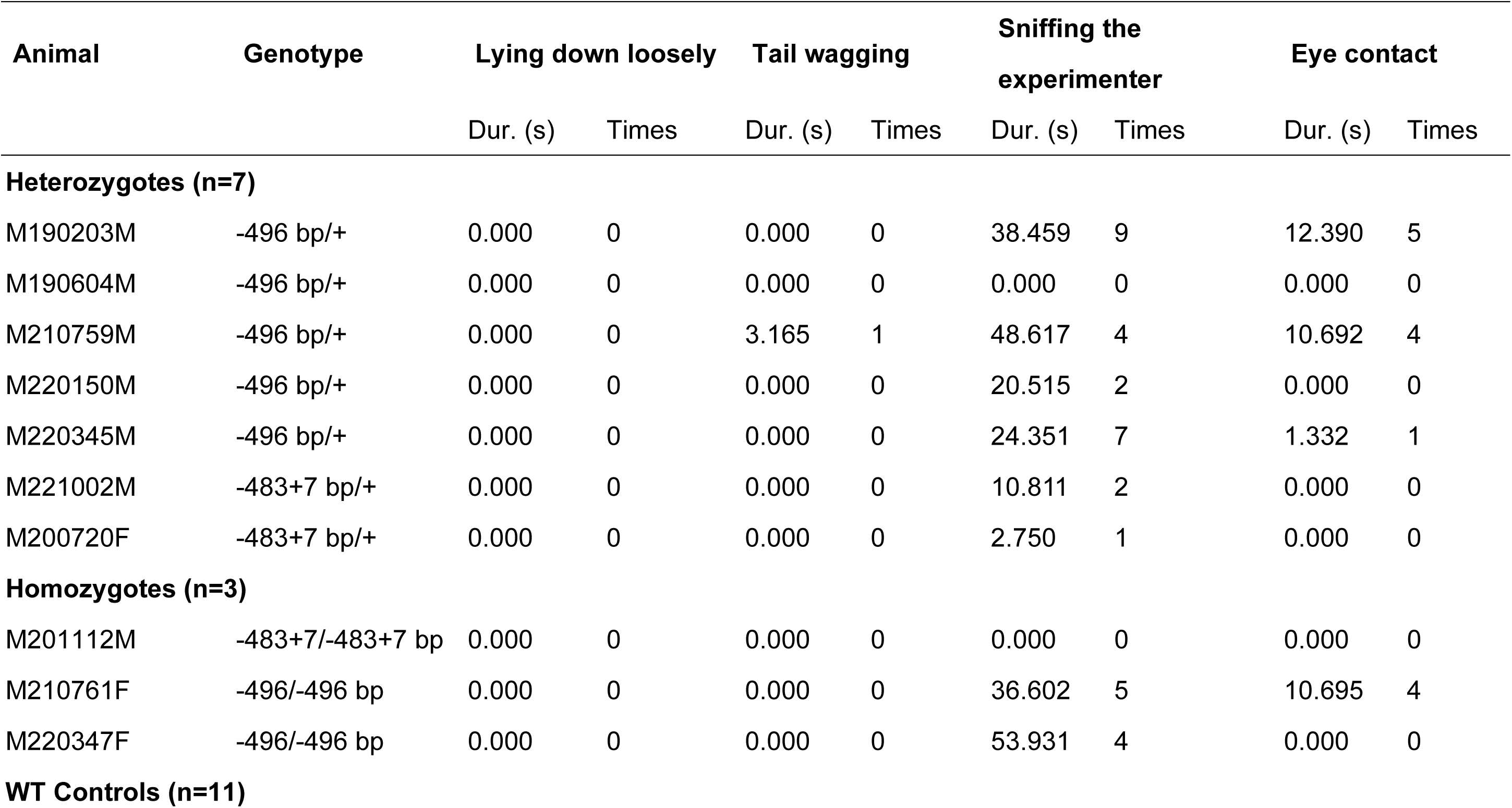
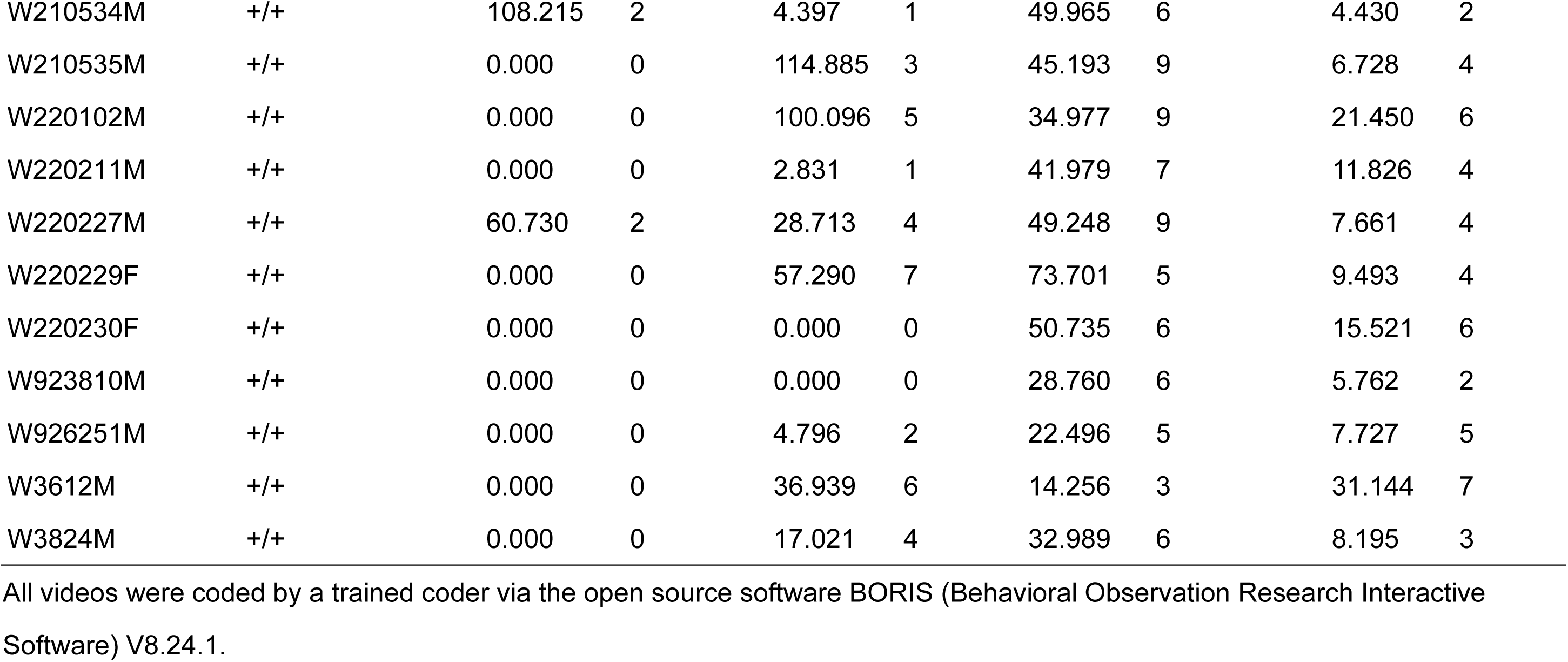
Comfort-related behaviors during phase 3 (hand stroke) of the affective touch assay.

**Supplementary Table 2.**
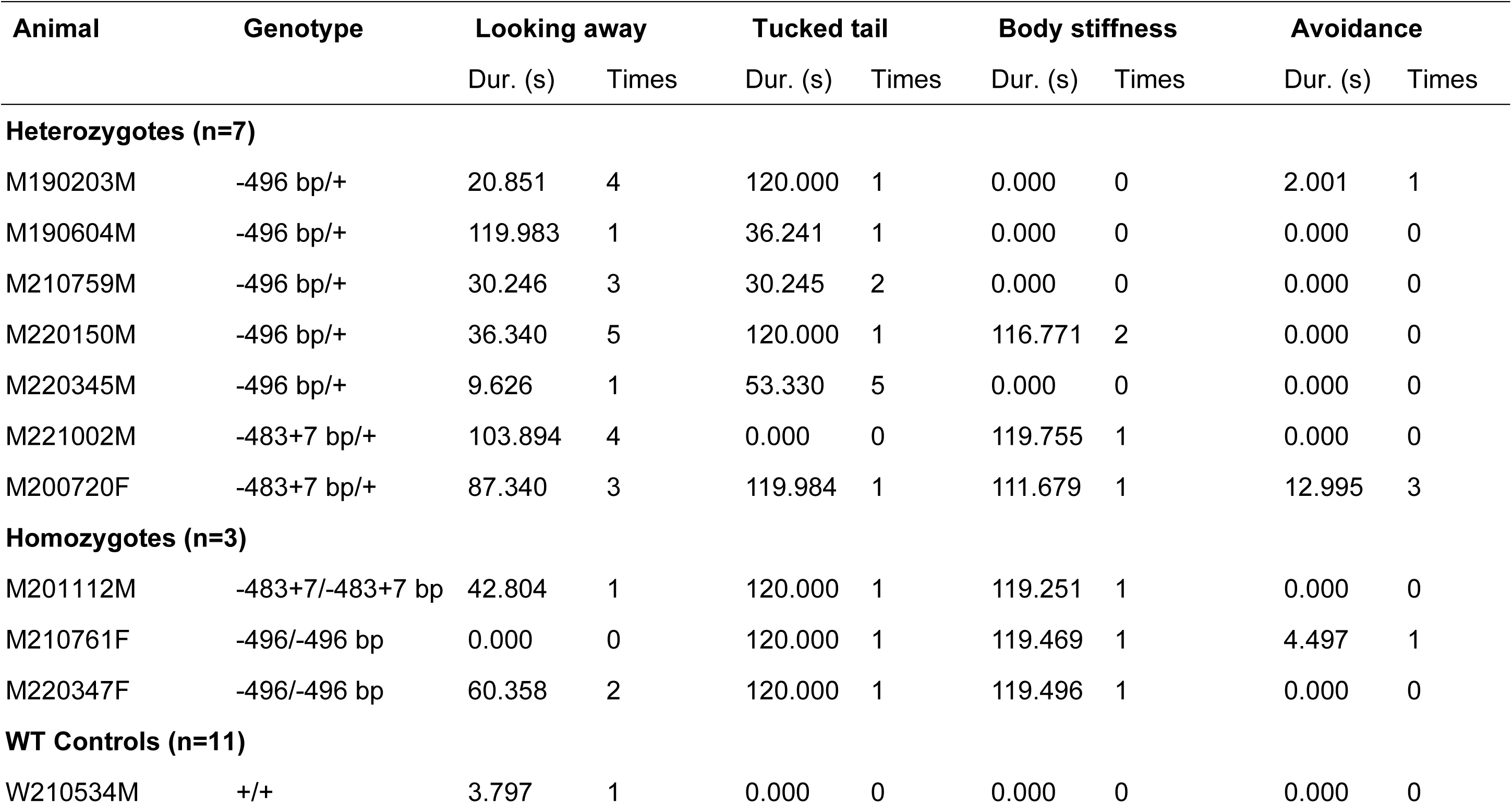
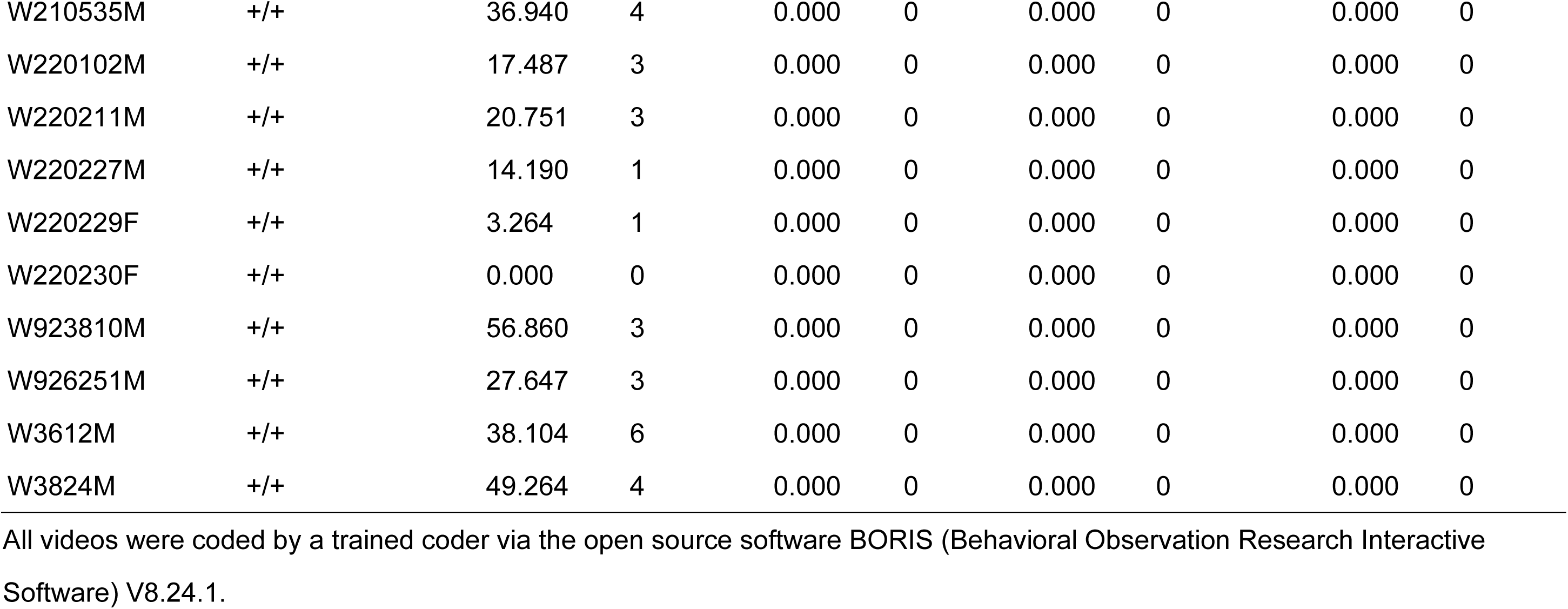
Stress-related behaviors during phase 3 (hand stroke) of the affective touch assay.

